# A New Method to Correct for Habitat Filtering in Microbial Correlation Networks

**DOI:** 10.1101/467365

**Authors:** Vanessa Brisson, Jennifer Schmidt, Trent R. Northen, John P. Vogel, Amélie Gaudin

## Abstract

Amplicon sequencing of 16S, ITS, and 18S regions of microbial genomes is a commonly used first step toward understanding microbial communities of interest for human health, agriculture, and the environment. Correlation network analysis is an emerging tool for investigating the interactions within these microbial communities. However, when data from different habitats (e.g sampling sites, host genotype, etc.) are combined into one analysis, habitat filtering (co-occurrence of microbes due to habitat sampled rather than biological interactions) can induce apparent correlations, resulting in a network dominated by habitat effects and masking correlations of biological interest. We developed an algorithm to correct for habitat filtering effects in microbial correlation network analysis in order to reveal the true underlying microbial correlations. This algorithm was tested on simulated data that was constructed to exhibit habitat filtering. Our algorithm significantly improved correlation detection accuracy for these data compared to Spearman and Pearson correlations. We then used our algorithm to analyze a real data set of 16S-V4 amplicon sequences that was expected to exhibit habitat filtering. Our algorithm was found to effectively reduce habitat effects, enabling the construction of consensus correlation networks from data sets combining multiple related sample habitats.

## 1 Introduction

The importance of microbial communities has become increasingly recognized in a variety of contexts, including human health, agricultural sustainability, and the conservation of our natural environment and resources. While many studies have evaluated microbial community composition, generally through amplicon (16S, ITS, 18s) based sequencing approaches, an increasing number of studies address community structure and interactions through microbial correlation network analyses. Microbial correlation networks have been used to investigate microbial communities in a wide range of systems including oceans, human microbiomes, and soil and plant associated microbial communities (Barberán et al., 2012; Delgado-Baquerizo et al., 2018; Faust et al., 2012; Lima-Mendez et al., 2015; Milici et al., 2016; Shi et al., 2016).

Microbial correlation network analysis goes beyond simple surveys of community composition, and can provide important insights into the ecological interactions within microbial communities. These analyses can be used to investigate the interactive structure of microbial communities across different experimental conditions and how communities change over time. For instance, a study of bulk soil and rhizosphere microbiomes from wild oat (*Avena fatua*) found greater complexity in the rhizosphere correlation networks as compared to bulk soil, with increasing network complexity over the course of plant growth that was repeatable over two growth cycles (Shi et al., 2016). Increasing network complexity has been interpreted to indicate increased ecological interactions such as mutualistic metabolic cross-feeding and niche sharing (Berry and Widder, 2014; Shi et al., 2016). Microbial correlation networks have also been used to identify keystone species that appear to play a central role in microbial community structure. For instance, Shi et al. evaluated within and between module connectivity of network nodes to try to identify hub species (Shi et al., 2016), while Berry and Widder found that high degree centrality, high closeness centrality, and low betweenness centrality were indicative of keystone species in microbial correlation networks (Berry and Widder, 2014). Correlation analyses have also been used to generate testable hypotheses about specific interactions. In a study of plankton communities, an interaction between acoel flatworms (*Symsagittifera* sp.) and a green microalgae (*Tetraselmis* sp.) was predicted based on correlation analysis, and subsequently confirmed via microscopy (Lima-Mendez et al., 2015).

Several different approaches have been used to construct microbial correlation networks. While some studies use Spearman or Pearson correlations between microbial abundances, other tools have been introduced to address the particular issues associated with microbial community analysis (Deng et al., 2012; Faust and Raes, 2016; Friedman and Alm, 2012; Kurtz et al., 2015; Weiss et al., 2016). For instance, because the data represent relative abundances, these data are inherently compositional: an increase in the relative abundance of one taxon is necessarily accompanied by a decrease in the relative abundance of other taxa. Compositionality can also induce spurious correlations, a problem that is especially important in lower diversity systems (Friedman and Alm, 2012). Tools such as SparCC and SPEIK-EASI have been introduced specifically to address compositional effects in these analyses (Aitchison, 1982; Friedman and Alm, 2012; Knight et al., 2018; Kurtz et al., 2015). A comparison of network construction approaches details pros and cons of several of these approaches for detecting particular types of interactions (Weiss et al., 2016).

Habitat filtering (HF) can confound microbial correlation network analysis when samples from different habitats (e.g. sampling sites, soil compartments, host genotype) are combined (Berry and Widder, 2014; Röttjers and Faust, 2018). HF occurs when microorganisms’ abundances are correlated with different habitats. For example, two microorganisms that are correlated with habitat will appear to be correlated with each other when data from different habitats are combined into one analysis, as conceptualized in Figure 1. The two microorganisms’ abundances are not correlated in Habitat A or in Habitat B, but both increase in abundance with the change in habitat between A and B (Figure 1-A, B). When data from Habitat A and Habitat B are combined, the two microorganisms appear to be strongly correlated (Figure 1-C). However, this correlation is the result of HF rather than an underlying interaction between the organisms. Correction for HF, as proposed here, eliminates detection of the spurious correlation (Figure 1-D).

**Figure 1.**
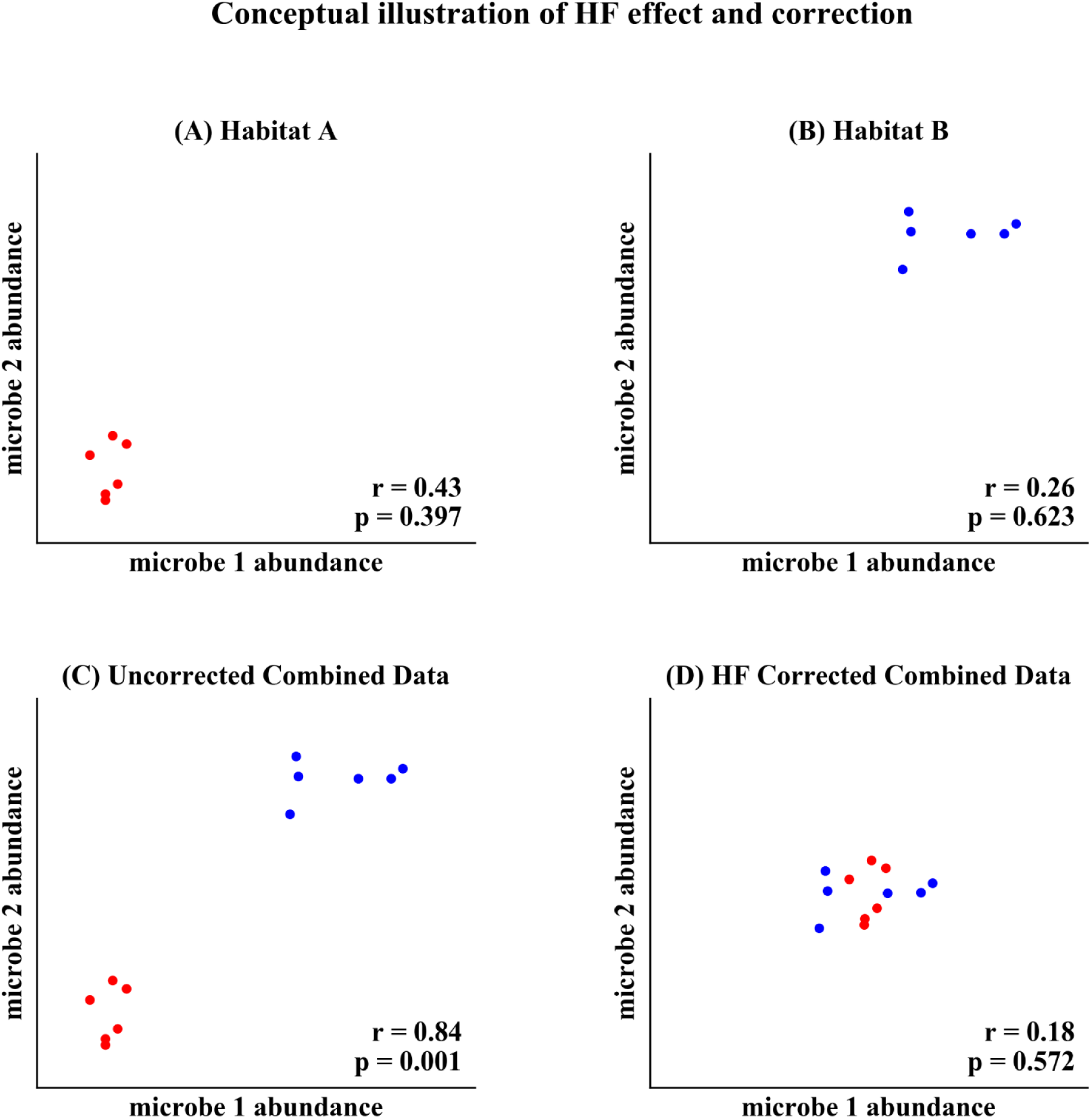
Conceptual illustration of HF effect and correction. (**A**) Two microorganisms’ abundances in Habitat A. (**B**) The same microorganisms’ abundances in Habitat B. (**C**) Combined data from Habitat A and Habitat B, resulting in a habitat induced correlation. (**D**) Correction for HF by subtracting the within habitat mean abundance eliminates detection of the spurious correlation.

Habitat effects have been shown to dominate network structures when data from different habitats are combined into a single analysis. For instance, an analysis of data from the human microbiome project found that the network clustered largely by sample site on the human body (Faust et al., 2012). Similarly, a recent study of soil bacteria across a range of habitats found that bacterial habitat preference dominated co-occurrence network structure (Delgado-Baquerizo et al., 2018).

Some studies have tried to identify habitat related effects in correlation network analyses, generally by addressing the impacts of specific measurable habitat factors. In a study of plankton communities, several factors including temperature, mixed layer depth, and phosphate and nitrite concentrations were shown to have significant effects on microbial abundances and induce network correlations (Lima-Mendez et al., 2015). In that study, the authors analyzed “taxon-taxon-environment associations” in which two taxa correlated with each other and with a particular environmental factor, and used that analysis to remove some spurious correlations from the network (Lima-Mendez et al., 2015). Another study proposes an algorithm to determine global interaction coefficients that account for gradients of environmental factors (Shang et al., 2017). However, both of these analyses were facilitated by relatively large sample sizes (313 samples and 150 samples respectively) and also did not account for additional environmental factors that were not measured experimentally but may be of importance for community composition and structure (Lima-Mendez et al., 2015; Shang et al., 2017). For example, plant root exudates are complex mixes of organic compounds that have been shown to influence microbial community composition in the rhizosphere (Zhalnina et al., 2018). Even closely related plant genotypes can differ in exudate composition, which may influence microbial community composition (Iannucci et al., 2017). Thus, failing to account for HF, due to plant genotype for instance, may bias networks with spurious correlations.

In this study we propose and test a new algorithm for HF correction that could be used for microbial network analysis across multiple study systems. The aims of this approach are (a) to enable the construction of accurate consensus networks from multiple habitats by removing habitat induced edges, (b) to do this with relatively small sample sizes, and (c) when differences between habitats are not necessarily easily characterized.

## 2 Results

### 2.1 An algorithm for HF correction

We developed an algorithm that corrects for HF effects in microbial abundance data prior to correlation detection, enabling more accurate and unbiased identification of the underlying microbial correlations (Figure 1-D). In these data sets, microorganism abundances are approximated by the (relative) abundances of amplicon sequence variants (ASVs; alternately operational taxonomic units or OTUs) in the sequencing data. The HF correction algorithm subtracts the within habitat mean abundance for each ASV from the abundances for that ASV in each sample. Thus, for a data matrix of abundance data A, the elements of the corresponding HF corrected matrix C are given by Equation 1.

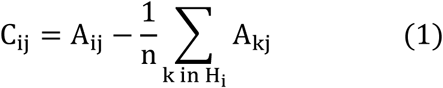

Here A_ij_ represents the abundance of ASV j in sample i, H_i_ represents the set of all samples for the habitat from which sample i originated, and n represents the number of samples in H_i_. Once correction has been performed, microbial correlations can be detected with either Spearman or Pearson correlations based on the corrected data matrix C. For the purposes of the analyses below, we have used Spearman correlations after HF correction.

In order to reduce correlations caused by the compositionality of relative abundance data, the data can be transformed to reduce compositionality effects prior to HF correction and correlation detection. For the analysis of real data described here, the centered log ratio transformation was used to account for compositionality (Aitchison, 1982).

### 2.2 HF correction improves accuracy of correlation detection for simulated data

We tested the performance of Pearson correlations, Spearman correlations, and the HF correction algorithm in detecting correlations in simulated data. Simulated data sets represented relative abundances of 50 ASVs, with each pair of ASVs having a 10% chance of having a true correlation, and each ASV having a 10% chance of being impacted by HF. Fifty simulations were evaluated at each of a range of sample sizes (6 to 60 samples = 3 to 30 samples per habitat) and HF strengths (0 to 10 times the ASV abundance standard deviation)

Two measures were used to evaluate algorithm performance on simulated data. First, we evaluated the root mean squared error (RMSE) of the detected correlation matrix as compared to the correlation matrix used in simulated data generation prior to the imposition of the HF effect.

This measure of correlation accuracy has been used previously to evaluate the performance of correlation detection algorithms to recover correlation patterns from simulated data (Friedman and Alm, 2012).

Second, the proportion of correlations detected that corresponded to true correlations in the simulated network was used as another evaluation of correlation accuracy. This was determined based on including all correlations detected with a significance cutoff of p < 0.01. These were compared to the randomly generated set of correlations from the initial stage of data simulation. Detected correlations that corresponded to true correlations with the same direction were considered correct, while those corresponding to no correlation or with the incorrect correlation direction were considered incorrect.

As compared to the other correlation detection methods tested (Spearman and Pearson), HF correction resulted in significantly lower RMSE and higher proportions of correlations correct for simulated data with HF (Figures 2-4). When no HF was imposed on the simulation (corresponding to 0 on the x axes of Figures 2, Figure 3, and Figure 4-A, B), all methods demonstrated similar performance. However, even a low level of HF (HF effect size equal to the standard deviation of the data) resulted in worse performance for Spearman and Pearson correlation networks, while the performance of HF correction remained stable. The difference in performance increased with increasing strength of the HF effect. When tested over a range of sample sizes and HF strengths, these effects were statistically significant (p < 0.05) for all combinations except when both sample size and HF strength were very low, as indicated by the red boxes in Figure 2 and Figure 3. With either larger sample size or larger HF effect size, performance with HF correction was significantly better than that of the other methods.

**Figure 2.**
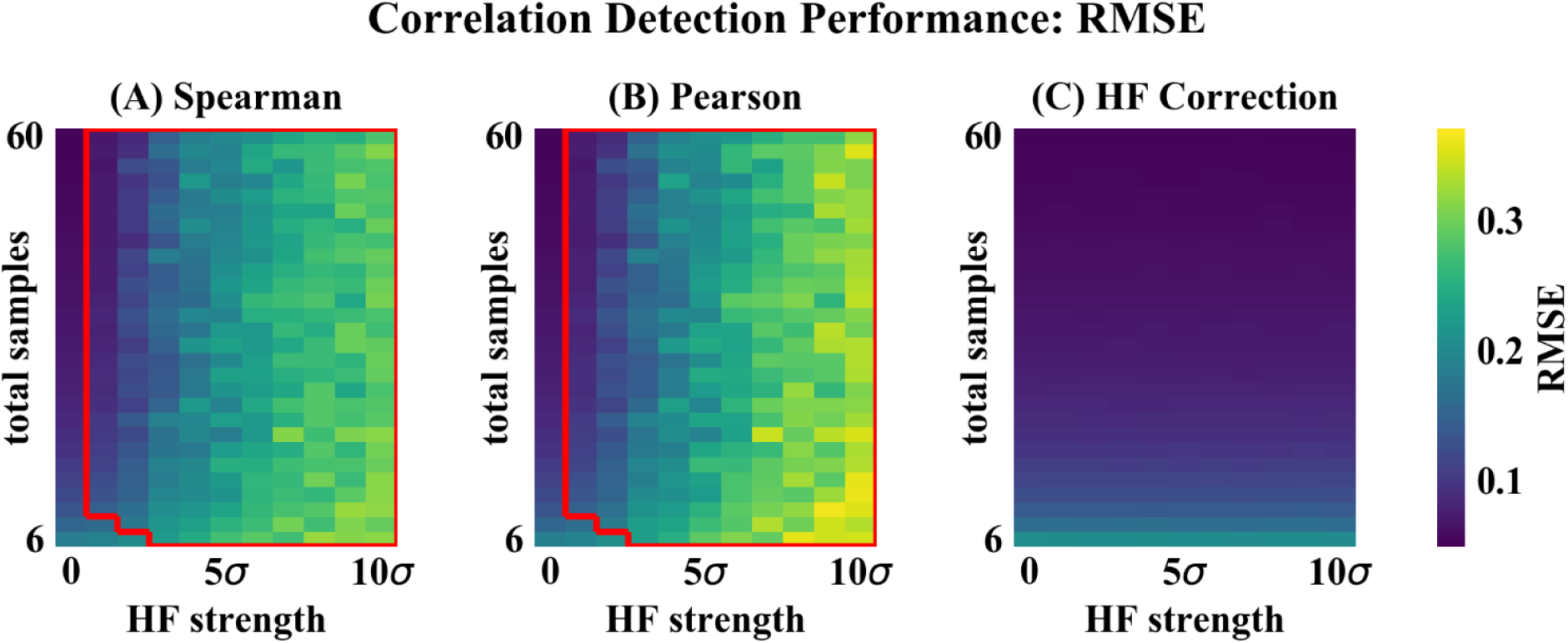
Correlation detection performance for simulated data: correlation matrix RMSE. Heat map colors represent the mean correlation matrix RMSE based on 50 simulated data sets for each combination of HF strength and sample size. (**A**) Spearman correlations. (**B**) Pearson correlations. (**C**) HF correction. Red box in (**A**) and (**B**) indicates the set of conditions (HF strength and sample size) for which there was a statistically significant improvement for HF correction compared to the other method tested.

**Figure 3.**
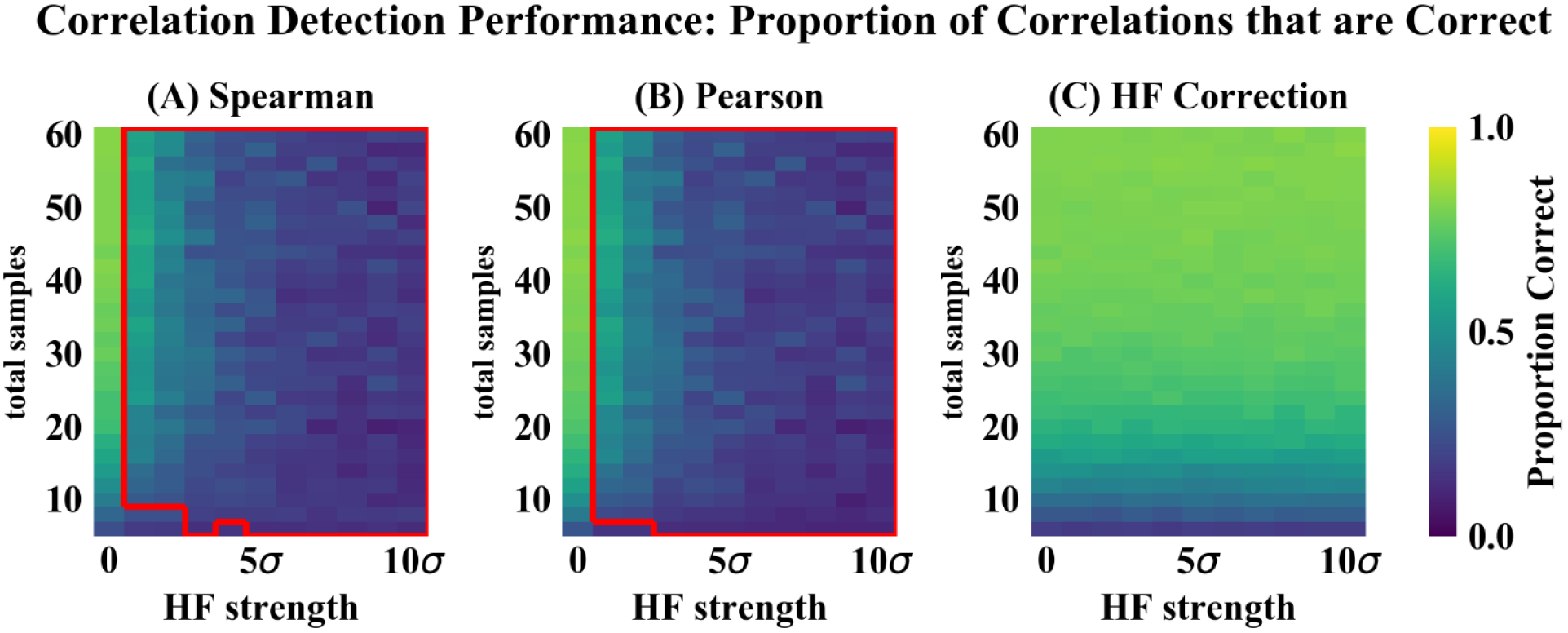
Correlation detection performance for simulated data: proportion of detected correlations that are correct. Heat map colors represent the mean proportion of correct correlations based on 50 simulated data sets for each combination of HF strength and sample size. (**A**) Spearman correlations. (**B**) Pearson correlations. (**C**) HF correction. Red box in (**A**) and (**B**) indicates the set of conditions (HF strength and sample size) for which there was a statistically significant improvement for HF correction compared to the other method tested.

**Figure 4.**
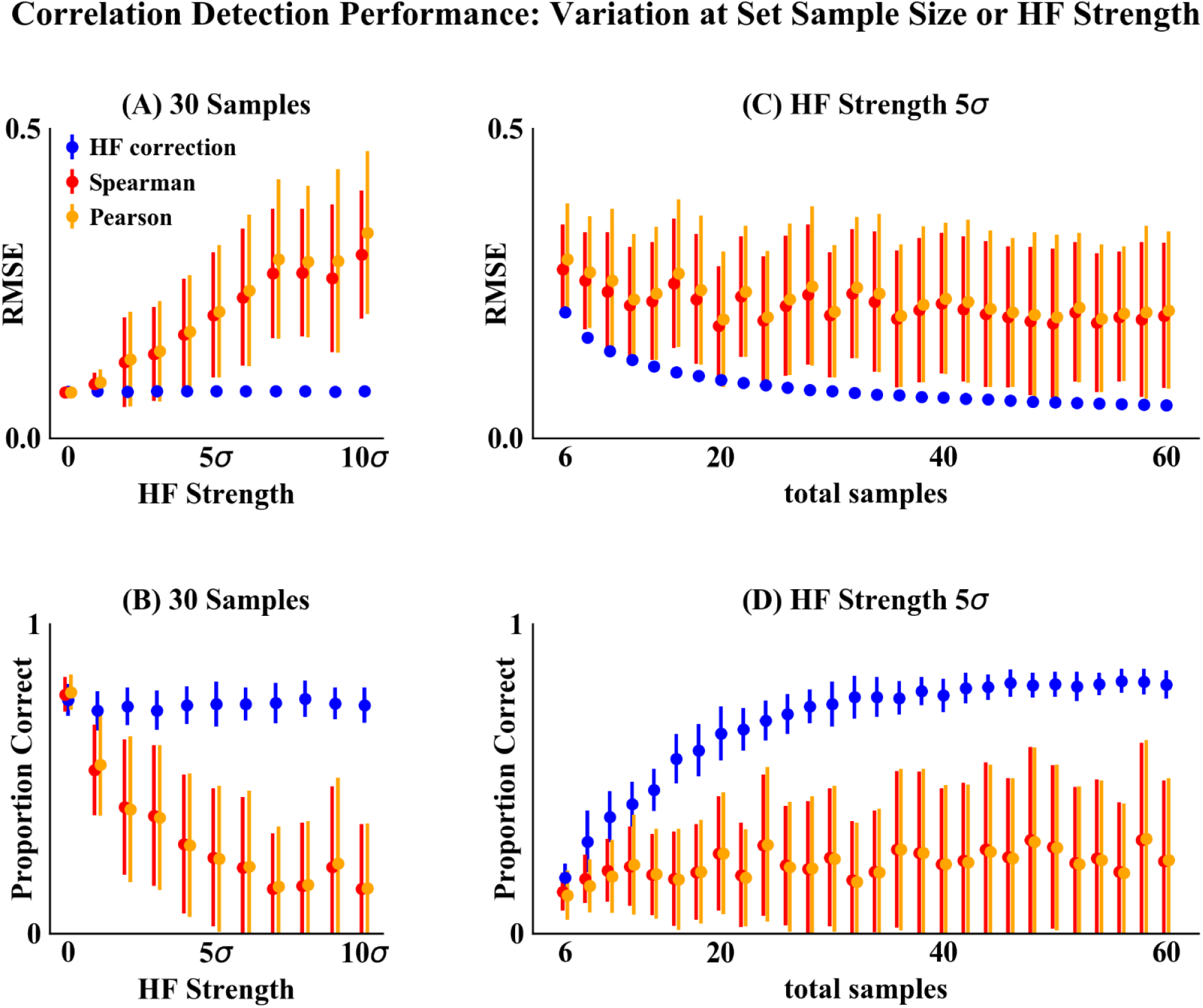
Correlation detection performance and variation on simulated data at set sample size or HF strength. (**A, B**) Set simulated sample size of 30 samples (15 samples for each habitat) and varying HF strength. (**C, D**) Set HF strength of 5σ and varying simulated sample size. Performance was measured based on two criteria: (**A, C**) RMSE between detected and simulated correlation matrices. (**B, D**) Proportion of correlations detected (at a significance cutoff of p < 0.01) corresponding to the initial randomly simulated correlations. Points indicate the mean and error bars indicate the standard deviation of 50 simulations.

### 2.3 HF correction reduces habitat preference bias in microbial correlation networks

In order to assess the HF correction algorithm’s performance on real microbiome data, we analyzed a data set expected to exhibit a significant HF effect: 16S-V4 amplicon sequences from rhizosphere soil collected from two distinct maize accessions grown in three differently managed soils. The two maize accessions were Mo17 (an inbred line that was sequenced to produce a maize reference genome) (Yang et al., 2017), and DeKalb2015 (a modern commercially successful transgenic hybrid line released in 2015). All three soils were from the Century Experiment at the Russell Ranch Sustainable Agriculture Facility (University of California, Davis), from replicated plots under different management practices and crop rotations for over 20 years (Wolf et al., 2018). The three soils’ histories are described in Table 1.

**Table 1.**
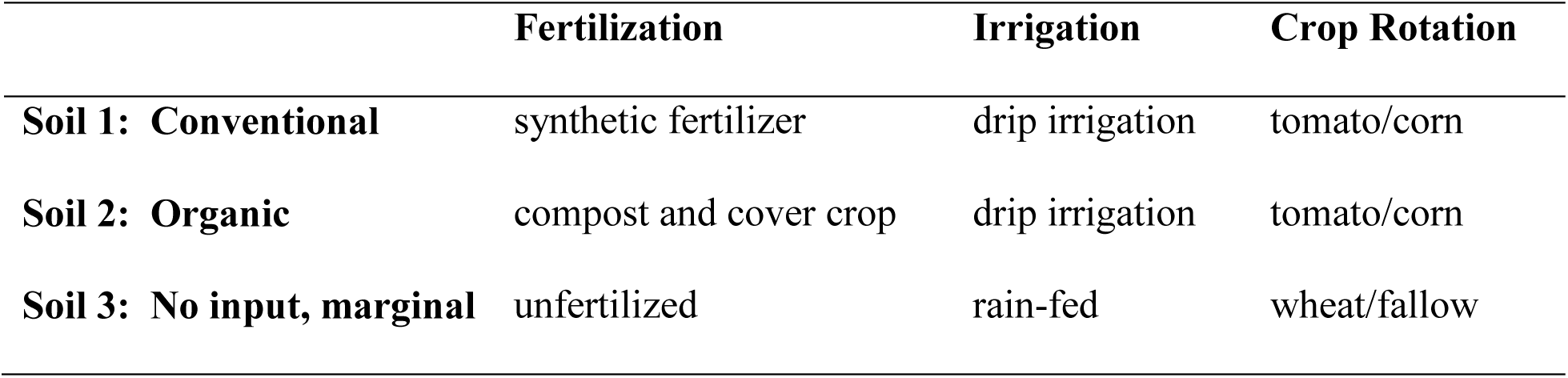
Soils used in experiment to explore rhizosphere microbial communities.

|  | **Fertilization** | **Irrigation** | **Crop Rotation** |
| --- | --- | --- | --- |
| **Soil 1: Conventional** | synthetic fertilizer | drip irrigation | tomato/corn |
| **Soil 2: Organic** | compost and cover crop | drip irrigation | tomato/corn |
| **Soil 3: No input, marginal** | unfertilized | rain-fed | wheat/fallow |

A principal coordinate analysis of the relative abundances of 16S-V4 ASVs from this experiment revealed a clear distinction between rhizosphere communities originating from two different maize accessions grown in the three different soils (Figure 5). There was a particularly clear difference between the communities from Soil 3 as compared to the other soils along the first principal coordinate axis. This analysis suggests a strong HF effect between the three soils.

**Figure 5.**
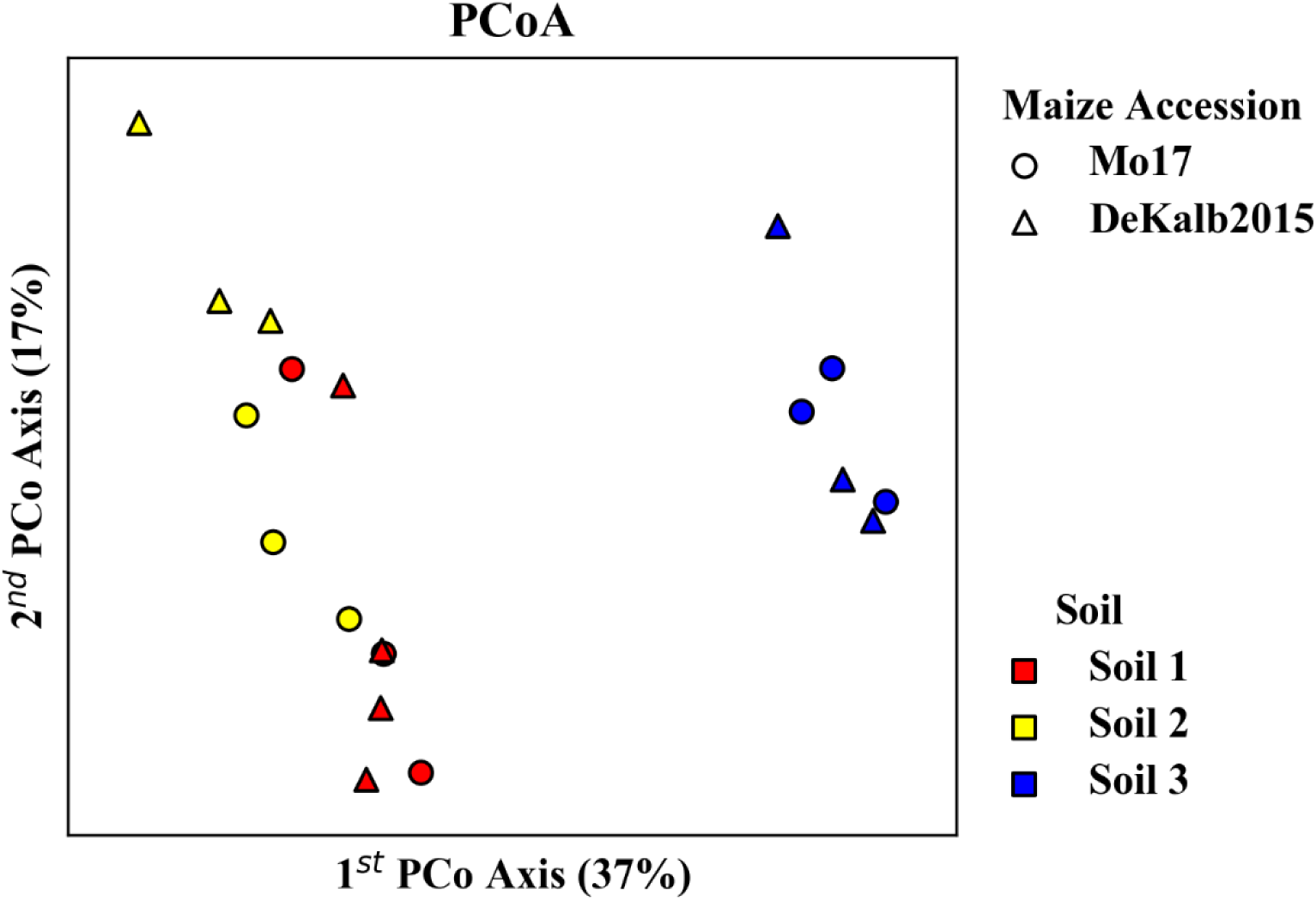
Principal coordinate analysis of rhizosphere microbiome data from two maize accessions grown in three soils. Analysis is based on 16S-V4 sequences using Bray-Curtis distances. The clear differences in community composition apparent between the different soils imply that HF occurs between soils.

Individual networks were constructed for each of the two maize accessions using Spearman correlations, Pearson correlations, and the proposed HF correction algorithm. Data were centered log ratio transformed prior to HF correction and/or correlation detection in order to account for inherent compositionality effects in microbiome data. Correlations were included in the network based on a maximum p-value of 0.01 and a minimum correlation strength of 0.75.

An independent differential abundance analysis was conducted on the same data to identify the habitat preference of individual ASVs in the microbiome communities (see Methods section for details of this analysis). Any ASVs that had a significantly (p_adjusted_ ≤ 0.05) increased abundance in samples from one (or two) of the soils as compared to the other(s) were considered to have a preference for the habitat(s). This information was used to evaluate how strongly network structures were influenced by HF effects.

The networks constructed with Spearman or Pearson correlations for both maize accessions were dominated by HF effects, while those constructed with HF correction exhibited a clear reduction or elimination of these effects (Figure 6). ASVs with the same habitat preference as determined by the independent differential abundance analysis (indicated by node color in Figure 6) clustered together in the Spearman and Pearson correlation networks. This indicates that HF effects rather than underlying microbial interactions dominate the structure of these networks. However, the networks constructed with the HF correction algorithm do not exhibit these patterns.

**Figure 6.**
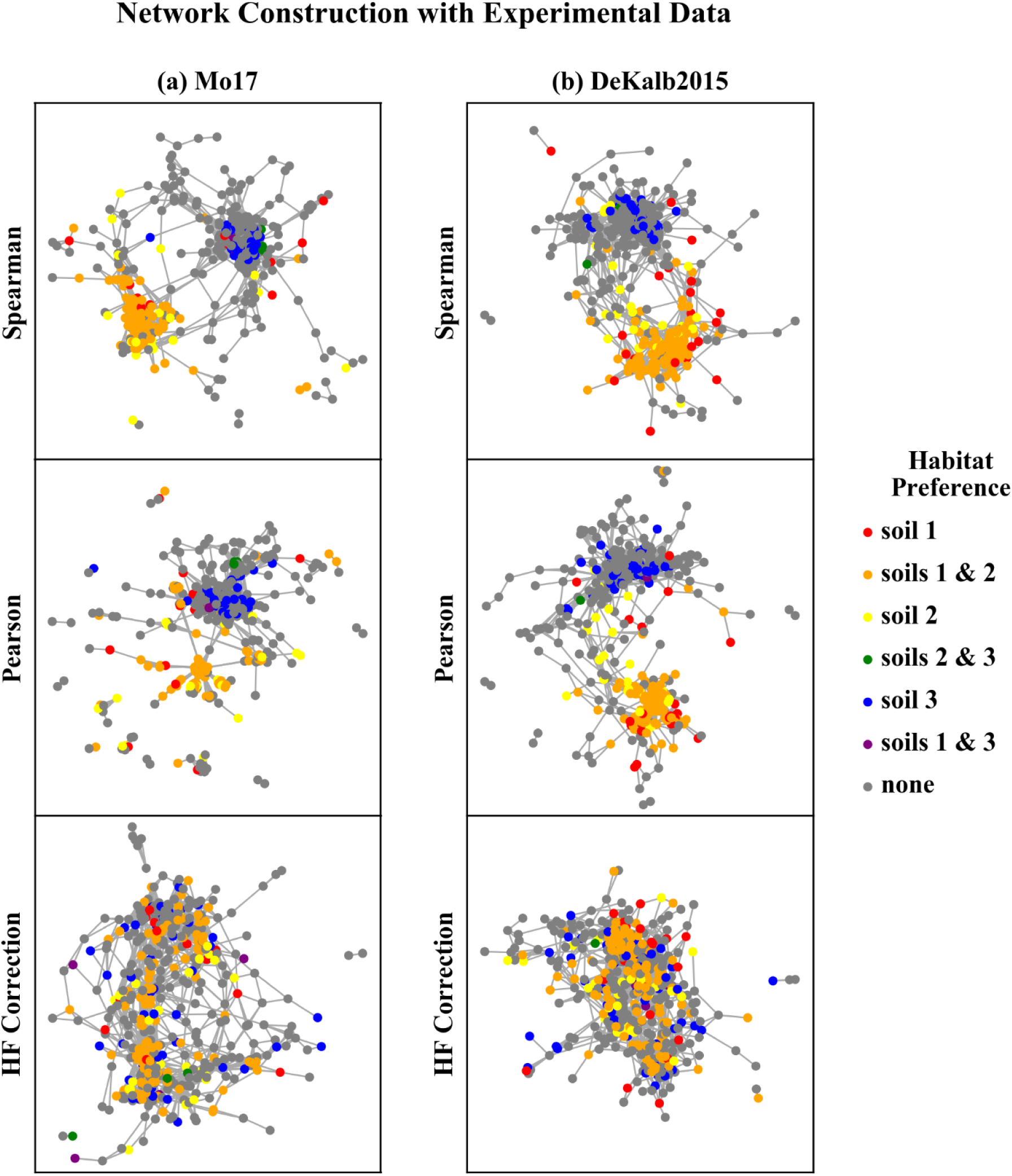
Rhizosphere microbiome correlation networks constructed with different correlation detection algorithms. Each node in the network represents a single ASV. ASVs are colored based on their habitat preference as assessed by the independent differential abundance analysis.

We quantified the extent to which the network structure corresponds to habitat preference using the proportion of network connections that are between ASVs with a shared habitat preference (Figure 7). For networks constructed for both maize accessions (Mo17 and DeKalb2015), the Pearson correlation networks had the highest proportion of correlations between ASVs with shared habitat preference (47% and 45%). Spearman correlation networks were intermediate (29% and 23%), and HF corrected networks had the lowest levels of shared habitat preference correlations (18% and 16%). The differences in habitat preference correlations between network construction algorithms were statistically significant (p < 0.01 based on chi-square test for equality of proportions). As a point of reference, for a randomized network based on this data set, this proportion would be 7% to 8%.

**Figure 7.**
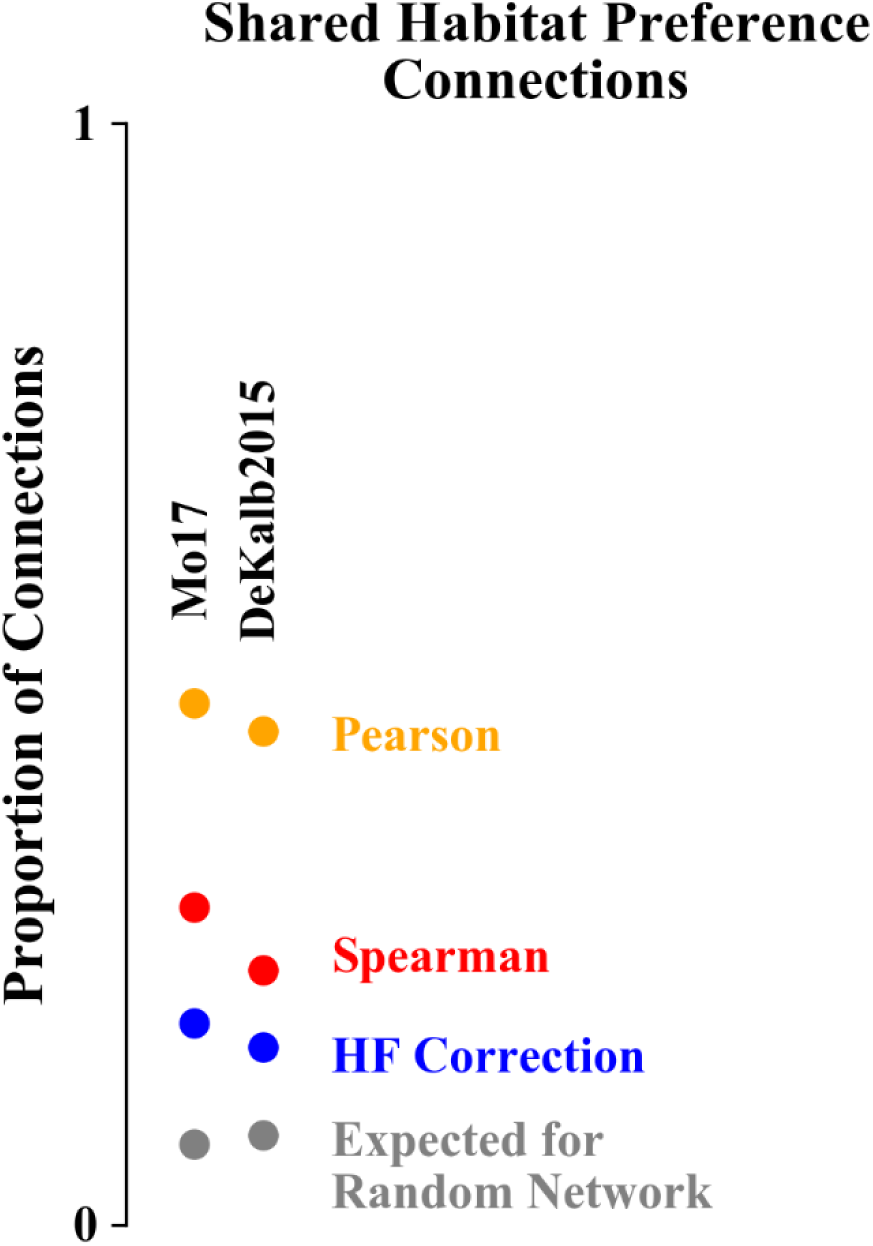
Proportion of network correlations that are between ASVs with a shared habitat preference. Networks constructed with Pearson correlations have the highest proportion of shared habitat preference correlations while HF corrected networks have the lowest.

## 3 Discussion

Correlation network analysis is an important tool for understanding interactions within microbial communities, but HF can have significant impacts that should be considered when constructing, analyzing, and interpreting these networks. HF effects are known to influence correlation network structure, and have been shown to affect a range of commonly used network construction approaches (Berry and Widder, 2014; Röttjers and Faust, 2018). In the analysis of rhizosphere microbiome sequencing data presented here, the uncorrected networks showed a strong impact of HF on the network structure (Figure 6 and Figure 7). This is consistent with the expectations based on both the soil history (Table 1) and the PCoA of the microbial community composition (Figure 5). Both crop rotation and fertilization practices are expected to influence the bulk soil microbial community, which in turn impacts the community that plants recruit to the rhizosphere from that soil. Another important factor affecting rhizosphere microbiomes is nutrient and water availability, which are strongly affected by soil management practices, and is also expected to influence rhizosphere microbiome recruitment by the plant (Koyama et al., 2014; Naylor et al., 2017).

We developed and tested an algorithm to correct for HF effects in correlation network construction when samples from different habitats are combined. The algorithm accomplishes the correction by subtracting the within habitat mean abundance from each ASV within each sample (Equation 1). This is conceptually similar to hierarchical regression, which has been used to analyze wood decay fungi co-occurrence (presence/absence) data to account for host tree preference when analyzing positive or negative associations between fungi (Ovaskainen et al., 2010).

The analysis of performance on simulated data showed that when HF was present, the HF correction was able to improve correlation detection compared to Spearman or Pearson correlations even at small sample sizes (n = 6 total samples (3 per habitat) for most HF strengths tested) (Figure 2 and Figure 3). Performance, measured as low RMSE and a high proportion of correct correlations, increased with increasing sample size up to approximately 30 total samples (15 per habitat), at which point performance leveled off (Figures 1-3). This is consistent with another study which recommended a sample size of at least 25 samples based on reduced specificity of network correlations at lower sample sizes (Berry and Widder, 2014).

Increasing HF strength negatively impacted the performance of Spearman and Pearson correlations for network construction. However, HF correction was able to maintain consistent performance, and lower variability in performance, across the full range of HF strengths tested. A recent comparison of several microbial correlation network construction methods, including CoNet (Faust and Raes, 2016), SparCC (Friedman and Alm, 2012), SPEIC-EASI (Kurtz et al., 2015), and Spearman correlations, also found that all methods tested exhibited reduced precision on data with simulated HF (Röttjers and Faust, 2018).

Understanding and accounting for HF effects is an important part of conducting meaningful microbial correlation network analyses of data from varying habitats. HF correction accounts for those effects and allows for the construction of consensus networks representing the underlying microbial correlations from data spanning multiple related habitats.

## 4 Methods

### 4.1 Simulated data generation

Data simulation was based on that described by Friedman et al., with some modifications (Friedman and Alm, 2012). Simulated data were generated for 50 ASVs, at a range of HF strengths (0 to 10σ) and a range of sample sizes (6 to 60 samples), with half of the samples (3 to 30) belonging to each of 2 simulated habitats. To generate an initial set of correlations, any pair of ASVs was given a 10% chance of being perfectly correlated, with equal probabilities that correlations were positive or negative. Based on those correlations, the nearest valid covariance matrix was determined based on a standard deviation of 0.1 using the cov_nearest function in the StatsModels Python module. Additionally, each ASV was given a 10% chance of exhibiting HF, with equal probabilities of positive or negative correlation with habitat. Data were drawn from a lognormal distribution with a standard deviation of 0.1. Fifty simulated data sets were generated for each combination of HF strength and sample size.

### 4.2 Experimental data collection and analysis

Experimental data were from rhizosphere soil samples from two different maize accessions grown in three different soil types as described above. Maize seeds were surface sterilized in 5% sodium hypochlorite for one minute, germinated in petri dishes, and planted in five gallon pots containing one of the three soils. Plants were grown in a greenhouse for eight weeks with automated drip irrigation and no fertilization. At the end of the experiment, plants were harvested and rhizosphere soil was collected as described by Barillot et al. (Barillot et al., 2013). DNA was extracted with the MoBio PowerSoil extraction kit [MoBio Laboratories Inc., Carlsbad, CA, USA] according to the manufacturer’s instructions.

16S-V4 amplicon sequencing was conducted at the DOE Joint Genome Institute using the 515F (GTGYCAGCMGCCGCGGTAA), 805R (GGACTACNVGGGTWTCTAAT) primer set. Sequencing was performed on an Illumina MySeq in 2×300 run mode. Sequencing data were analyzed with the DADA2 package in R based on the big data analysis pipeline with the following parameters: truncLen = (160, 120), maxEE = (2, 5), and truncQ = 2 (Callahan et al., 2016). The resulting sequence count data were analyzed using the phyloseq package in R (McMurdie and Holmes, 2013). Chloroplast and mitochondrial sequences were removed and data were rarefied to the minimum number of reads in all samples prior to further analysis.

### 4.3 Network Construction

Networks were constructed with each of three different approaches: Spearman correlations, Pearson correlations, and our proposed HF correction algorithm. For HF correction networks, rhizosphere microbial data were corrected based on equation 1 and correlation were determined from corrected data using Spearman correlations. For analysis of simulated networks, both positive and negative correlations were included. In order to select significant correlations to determine the proportion of correct correlations, a significance cutoff of p < 0.01 was applied. For analysis of experimental data, only ASVs detected in at least 50% of the samples being analyzed were included in the analysis. Experimental data were first transformed with the centered log transform in order to account for compositionality of the data. In addition to the significance cutoff of p < 0.01, a correlation strength cutoff of r > 0.75 was also applied and only positive correlations were included in the networks based on experimental data.

### 4.4 Differential abundance analysis

An independent differential abundance analysis was conducted to identify the habitat preference of individual ASVs. This analysis was conducted using DeSeq2 to compare ASV abundances between pairs of soil types (Love et al., 2014). P-values were adjusted to account for multiple comparisons using the Benjamini and Hochberg correction. Adjusted p-values below 0.05 were considered to indicate habitat preference for the associated ASV.

### 4.5 Data Availability

The DNA sequencing dataset used in this study is available through the JGI Genome Portal at https://genome.jgi.doe.gov/portal with the project ID 1138452. An implementation of the HF correction algorithm as a python module, along with the ASV abundance table analyzed in this study, is available through GitHub at https://github.com/vbrisson/HabitatCorrectedNetwork.

## Author Contributions

VB developed and implemented the habitat filtering correction algorithm. VB and JS tested the algorithm. AG and JS designed the rhizosphere experiment. JS performed the rhizosphere experiment. VB analyzed the results. TN, JV, and AG provided scientific advice. VB, JS, JPV, TN, and AG contributed to the discussion of the results and writing of the manuscript.

## Acknowledgments

The authors wish to thank the Community Science Program from the Joint Genome Institute for providing sequencing resources to AG and collaborators at Dupont Pioneer for sharing seeds from the ERA panel.

